# Insulin signal transduction is impaired in the type 2 diabetic retina

**DOI:** 10.1101/856245

**Authors:** Youde Jiang, Li Liu, Hainan Li, Jie-Mei Wang, Jena J. Steinle

## Abstract

Rates of type 2 diabetes are reaching epidemic levels. Yet, the tissue specific alterations due to insulin resistance are only recently being investigated. The goal of the present study was to evaluate retinal insulin signal transduction in a common mouse model of type 2 diabetes, the db/db mouse. Retinal lysates from five month old male db/db and db/+ (control) mice were collected and processed for Western blotting or ELISA analyses for insulin receptor, insulin receptor substrate-1 (IRS-1), Akt, tumor necrosis factor alpha (TNFα) and caspase 3 levels. Data demonstrate increased TNFα and IRS-1 phosphorylation on serine 307. This led to decreased Akt phosphorylation on serine 473 and increased cleavage of caspase 3. Taken together, the data suggest dysfunctional insulin signaling in the retina of the db/db mouse.

## Introduction

With increasing rates of obesity, rates of type 2 diabetes and diabetic complications are expected to rise exponentially over the next few decades (American Diabetic Association). A key feature of type 2 diabetes is a resistance to insulin. Insulin signaling is key to a number of physiological processes, including glucose metabolism, cell growth, general gene expression, and apoptosis. Studies have focused on insulin resistance in the insulin-responsive tissues with less focus on other organs, such as the retina. We have previously reported that diabetes-induced increases in TNFα can cause phosphorylation of insulin receptor substrate 1 (IRS-1) in Serine 307, thus inhibiting normal insulin signal transduction in retinal endothelial cells [1]. This increase in TNFα was also associated with increased cleavage of caspase 3. We found similar findings in BBZDR/Wor type 2 diabetic rats [2, 3]. However, it was not clear if these findings also occurred in type 2 diabetic mouse models.

For these studies, we used the db/db model of type 2 diabetes. We chose to use these mice as others have reported significant retinal changes. Work showed that intermittent fasting altered the gut microbiome in the db/db mice, which was associated with less retinal damage [4]. Additional studies also showed that diabetes in the db/db mice led to reduced diurnal oscillatory rhythms, which altered metabolic pathways [5]. Other groups reported increased permeability and inflammatory mediators in the retina of db/db mice, which was reduced by C1q/tumor necrosis factor (TNF) related protein 9 [6]. Ginsenoside Rg1 was shown to reduce retinal neurodegeneration in the db/db mouse through activation of IRS-1/Akt/GSK3β levels [7]. Since it is clear that the retinas of db/db mice have damage, we wanted ascertain whether this was due to altered insulin signal transduction.

We hypothesized that retinal lysates from db/db mice would have increased IRS-1^Ser307^ phosphorylation, leading to decreased Akt levels with increased cleavage of caspase 3.

## Methods

### Mice

Five month old male db/db (BKS.Cg-Dock7^m^+/+Lepr^db^, wildtype for Dock7^m^, homozygous for Lepr^db^) and db/+ (wildtype for Dock7^m^, wildtype for Lepr^db^, from the same colony) mice were used for these experiments. Mice were purchased from Jackson Laboratory (#000642) at 2 months and age and allowed to age to 5 months at the vivarium. All animal procedures meet the Association for Research in Vision and Ophthalmology requirements and were approved by the Institutional Animal Care and Use Committee of Wayne State University and conform to NIH guidelines. Animal body weights and glucose levels are in Table 1.

**Table 1.** Body weight (g) and blood glucose (mg/di) of db/+ (Control) and db/db mice at sacrifice. *P<0.05 vs. db/+. N=5.

|  | Body weight (g) | Blood glucose (mg/dl) |
| --- | --- | --- |
| Db/+ | 21.6 | 118 |
| Db/db | >60* | >600* |

#### Western blotting

Whole retinal lysates were collected into lysis buffer containing protease and phosphatase inhibitors. Equal amounts of protein were placed into pre-cast tris-glycine gels (Invitrogen, Carlsbad, CA), and blotted onto nitrocellulose membranes. After blocking in TBST (10mM Tris-HCl buffer, pH 8.0, 150 mM NaCl, 0.1% Tween 20) with 5% BSA, membranes were treated with a phosphorylated insulin receptor (Tyr 1150/1151), insulin receptor, phosphorylated Akt (Ser473), total Akt, phosphorylated insulin receptor substrate 1 (Ser307), total IRS-1 (Cell Signaling Technology, Danvers, MA) TNFα, (Abcam, Cambridge, MA), and beta actin (Santa Cruz Biotechnology, Santa Cruz, CA) primary antibodies overnight. The following day, membranes were incubated with secondary antibodies labeled with horseradish peroxidase. Antigen-antibody complexes were visualized using chemilluminescence (Thermo Scientific, Pittsburgh, PA). Data was analyzed on an Azure C500 machine (Azure Biosystems, Dublin, CA). Western blot band densities were measured using Image Studio Lite software.

### ELISA

A cleaved caspase 3 ELISA (Cell Signaling Technology, Danvers, MA) was done according to manufacturer’s instructions.

### Statistics

Data were assessed for changes db/+ control mice. Data are presented as mean ±SEM. P<0.05 was accepted as significant. Data was analyzed using Prism 8.0 (GraphPad software).

## Results

### Diabetes reduces insulin receptor and Akt phosphorylation

As we have shown in the retina from diabetic rats [3], diabetes significantly reduced insulin receptor and Akt phosphorylation in the db/db retina (Figure 1).

**Figure 1.**
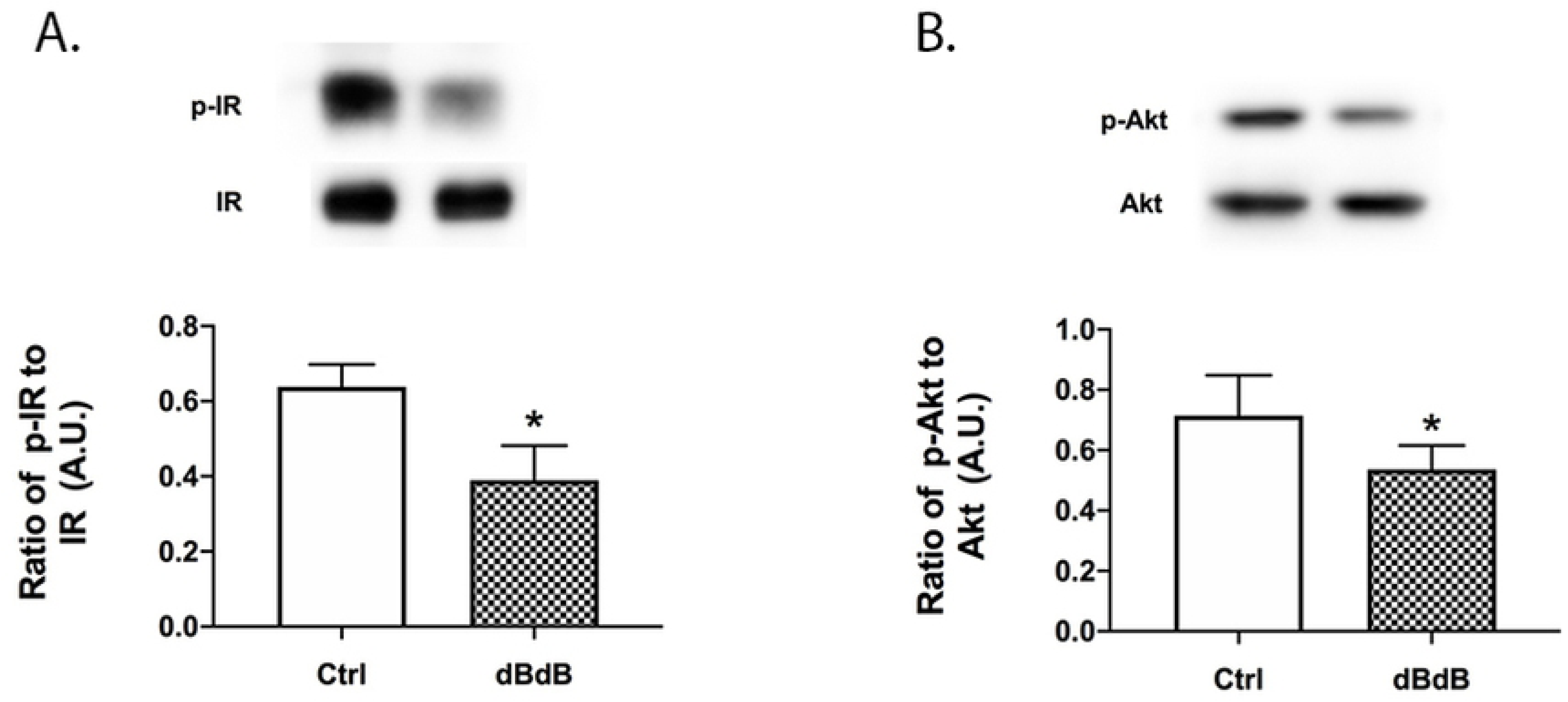
Western blotting for insulin receptor (A) and Akt phosphorylation (B) in db/+ (Control) and db/db mouse retinal lysates. *P<0.05 vs. Control. Data are mean ±SEM. N=5.

### TNFα and IRS-1^Ser307^ phosphorylation are increased in the db/db retina

With Akt phosphorylation reduced in the retina of db/db mice, one possible mechanism is due to increased IRS-1^Ser307^ phosphorylation. We have previously shown that diabetes-induced increases in TNFα can increase IRS-1^Ser307^ phosphorylation in retinal cell types [1, 8], which results in dysfunctional insulin signal transduction. Figure 2 shows increased TNFα and IRS-1^Ser307^ phosphorylation in the retina of db/db mice.

**Figure 2.**
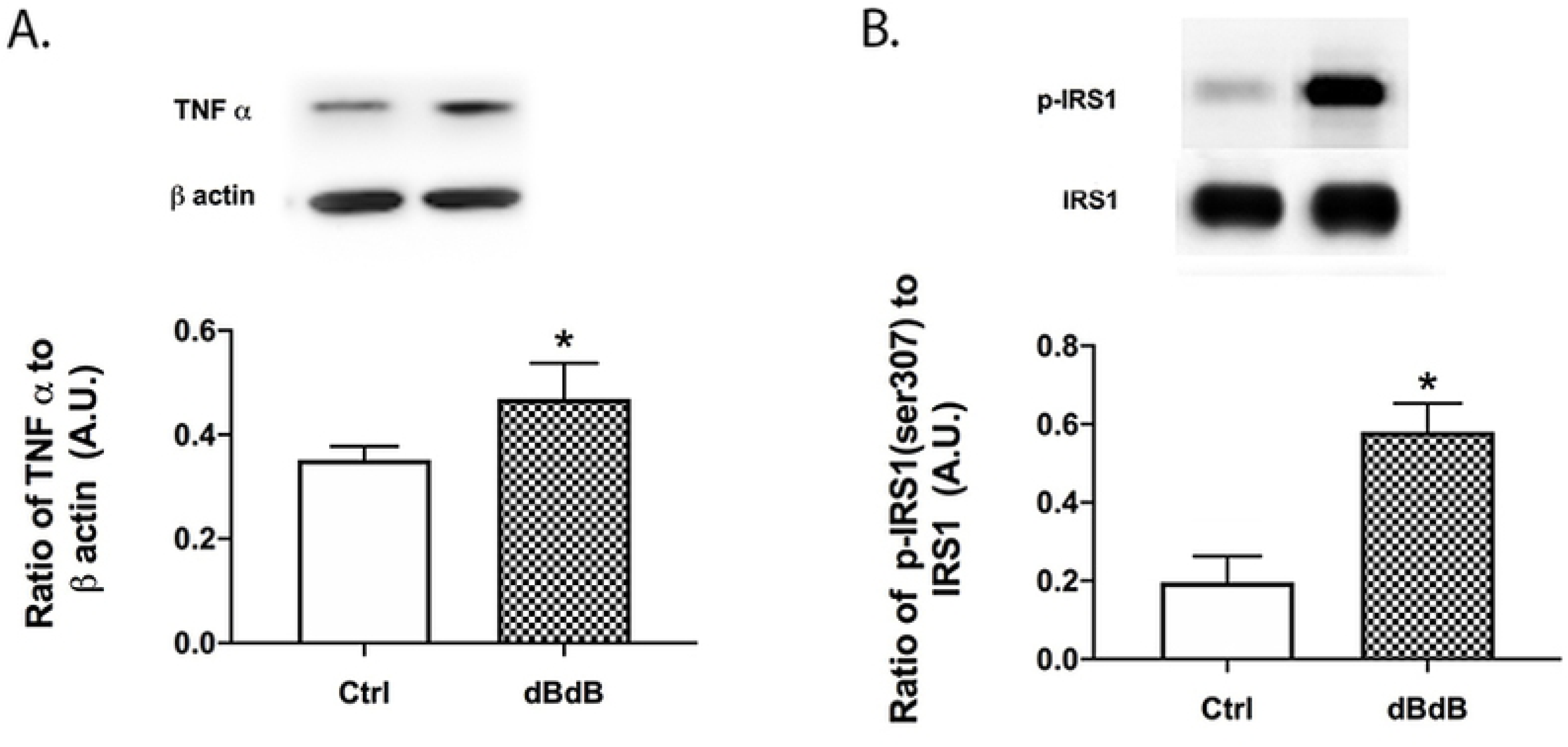
Western blotting for TNFα (A) and IRS-1^Ser307^ phosphorylation in db/+ (Control) and db/db mouse retinal lysates. *P<0.05 vs. Control. Data are mean ±SEM. N=5.

### Cleaved caspase 3 is increased in the type 2 diabetic retina

With reduced Akt phosphorylation, it follows that cleaved caspase 3 levels are significantly increased in the retina of the db/db mice (Figure 3).

**Figure 3.**
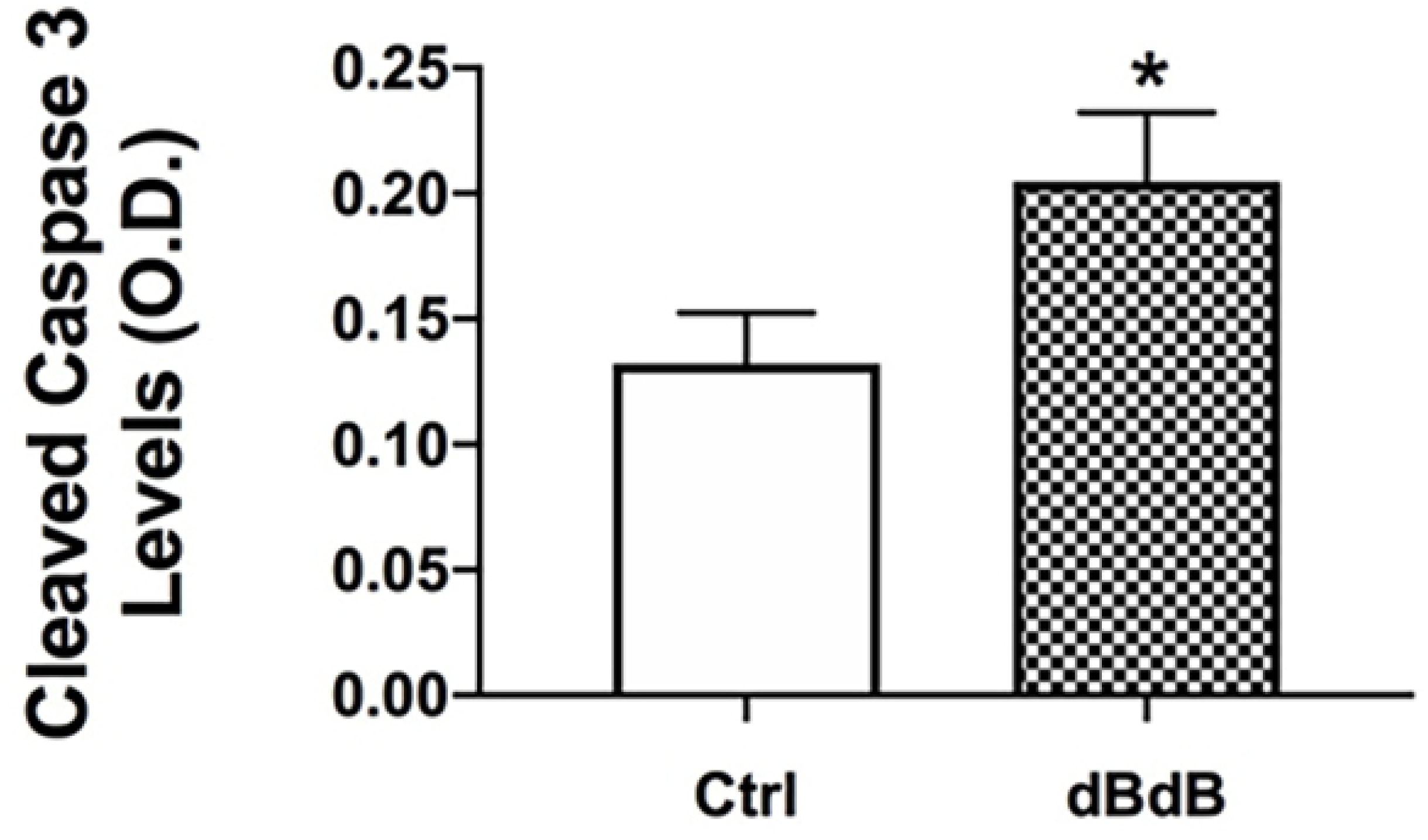
ELISA results for cleaved caspase 3 in db/+ (Control) and db/db mouse retinal lysates. *P<0.05 vs. Control. Data are mean ±SEM. N=5.

## Discussion

Type 2 diabetes is characterized by insulin resistance. However, this typically applies to insulin-responsive tissues, such as muscle, adipose tissue, and the liver. We have previously reported altered insulin receptor signaling in toll-like receptor 4 knockout mice [9], miR15a mice [10], and in retinal endothelial cells [1] and Müller cells [8]. However, none of these models mimic type 2 diabetes. We have shown retina damage in the type 2 diabetic rat model, the BBZDR/Wor rat [2, 3]. In this study, we wanted to investigate insulin signal transduction in the retina of db/db mice.

Studies on jejunal proteins from db/db mice showed impaired muscle insulin signaling, leading to insulin resistance [11]. Work has shown that rexinoids improved insulin signaling in skeletal muscle through decreased IRS-1^Ser307^ phosphorylation [12]. In contrast, work in hepatic tissues suggest that protein kinase C delta (PKCd) alters liver insulin signaling [13]. Focusing on the retina, studies have shown altered insulin signaling in the STZ model of type 1 diabetes [14]. In the STZ model, PKC altered insulin receptor immunoreactivity and signaling in endothelial cells and pericytes [14]. Additionally, work in the STZ model showed that insulin receptor signaling is key to health of the retinal pigmented epithelial (RPE) cells, leading to proper photoreceptor function [15].

Our findings in the present study suggest that retinal insulin signaling in impaired in the db/db mouse. The findings of increased TNFα levels associated with increased IRS-1^Ser307^ phosphorylation suggest that this may be the causative factor in the increased cleavage of caspase 3. It is established that the db/db mouse has altered retinal inflammation, which likely involves TNFα [6]. Increased TNFα can impair normal insulin signaling. Thus, retinal insulin signal transduction is impaired in the db/db mouse, similar to other models of diabetes.

